# Lignin degradation and valorization by *Pseudomonas putida* KT2440: a new role for glutathione peroxidase

**DOI:** 10.1101/2025.11.24.690113

**Authors:** Qing Zhou, Annabel Fransen, Paolo Innocenti, Arthur F.J. Ram, Johannes H. de Winde

## Abstract

Lignin, a complex natural aromatic polymer, poses significant challenges to its efficient degradation, hindering the utilization of biomass for many industrial applications. Bacterial degradation of lignin may offer a promising solution to this challenge. This project aimed at elucidating the function of secreted oxidative enzymes from *Pseudomonas putida* involved in lignin degradation and utilization. Using CRISPR-Cas9 and CRISPR-Cas3 systems, the putative lignin-degrading versatile peroxidase gene (VP; *PP*_*1686*, originally annotated as glutathione peroxidase GPx) and dye-decolorizing peroxidase gene (*PP_3248*) were individually knocked out from *P. putida* KT2440. The ΔPP_1686 mutant exhibited impaired growth and utilization of lignin-derived compounds. This correlated with reduced expression of p-hydroxybenzoate hydroxylase *pobA* and of DNA repair modules, alongside compensatory upregulation of energy and redox supply pathways. This work expands our knowledge on bacterial glutathione peroxidase by presenting a role beyond ROS scavenging. This work revealed the importance of *P. putida* VP/GPx in maintaining redox balance while supporting lignin-derived aromatic metabolism, offering new targets for future investigation into stress–metabolism crosstalk and lignin valorization strategies.

## 1. Introduction

Lignocellulose, the second most abundant terrestrial polymer on Earth, is composed of cellulose, hemicellulose, and lignin (Wang et al., 2019). Among those, lignin constitutes 15-40% of the dry weight of plant cell walls, making it the largest renewable source of aromatic biopolymers (Li et al., 2024; Ragauskas et al., 2014). Lignin is a highly complex aromatic macromolecule that is synthesized from phenylpropanoid precursors by polymerization in plants (Vogt, 2010). It is primarily derived from three monolignols—*p*-coumaryl alcohol, coniferyl alcohol, and sinapyl alcohol—which give rise to three major structural units in the lignin polymer: *p*-hydroxyphenyl (H), guaiacyl (G), and syringyl (S) moieties (Fu et al., 2021; Vanholme et al., 2010). The heterogeneous composition and diverse linkages between these building blocks result in a complex structure, making lignin highly recalcitrant and difficult to depolymerize and valorize. Therefore, current utilization of lignin is mainly limited to combustion for heat and energy production, and the use of raw lignin in relatively low-value products such as glues, resins, and asphalt (Demuner et al., 2019; Lora & Glasser, 2002; Stewart, 2008).

Towards lignin valorization, various chemical approaches have been explored. However, chemical degradation generates complex mixtures and undefined polymers that are unsuitable for value-added applications (Kleine et al., 2013; Smit et al., 2024; van Erven et al., 2024). Biological funneling strategies may provide new routes to overcome the challenges (Linger et al., 2014; Zhou et al., 2025). Recent studies have implicated several bacterial species capable of lignin degradation and utilization (Bugg, 2024; Weng et al., 2021), among which the Gram-negative soil bacterium *Pseudomonas putida* is considered one of the most promising (Xu et al., 2022). *P. putida* is known for its ability to convert a wide range of lignin-derived aromatic hydrocarbons, such as p-hydroxybenzoate, benzene, and xylene (Jiménez et al., 2002). Owing to these metabolic capacities, *P. putida* has already been engineered as a robust bacterial chassis (Nikel & de Lorenzo, 2018). For example, *P. putida* KT2440 was successfully engineered to metabolize lignin-derived compounds to valuable products like *cis, cis*-muconic acid (CCMA), polyhydroxyalkanoate (PHA) (Liu et al., 2024; Xu et al., 2021). Thus, *P. putida* has the potential to valorize lignin for future industrial applications. However, a clear enzymatic route from complex lignin towards well-defined lignin-derived compounds has not been clearly described.

In various microorganisms, typical oxidative enzymes, including laccase (Lac, EC 1.10.3.2), lignin peroxidase (LiP, EC 1.11.1.14), manganese peroxidase (MnP, EC 1.11.1.13), dye-decolorizing peroxidases (Dyp, EC 1.11.1.19), and versatile peroxidase (VP, EC 1.11.1.16), have been implicated in playing a role in the degradation of lignin (Pollegioni et al., 2015). Most of these enzymes were discovered in fungi. Bacterial enzymes implicated in lignin degradation have been much less studied, however, may provide more specific and controllable routes for aromatic compound transformation (Zhou et al., 2025). Several potentially ligninolytic enzymes, including laccase, Mn2+-independent peroxidases (e.g., DyPs), Mn2+-oxidizing peroxidases, multicopper oxidase (CopA), β-etherase, and dioxygenases were identified in the secretome of *P. putida* KT2440 (Xu et al., 2022). However, a specific function of these extracellular enzymes in lignin degradation was not further characterized. Hence, although suggestive, proof and understanding of lignin degrading enzymes in *P. putida* remained limited.

A recent study identified *P. putida* NX-1, isolated from leaf mold samples, as being capable of lignin degradation and able to grow on Kraft lignin as a sole carbon source (Xu et al., 2018). Degradation of lignin was determined using absorption at 280 nm (Chai et al., 2014). Genome analysis of *P. putida* NX-1 indicated the presence of putative enzymes involved in lignin decomposition, including Dyp-type peroxidases, versatile peroxidases (VP), manganese peroxidases (MP), and laccases (Xu et al., 2021), but a role in lignin degradation was not further substantiated.

Extracellular oxidative degradation of lignin is predicted to release reactive oxygen species (ROS). Glutathione peroxidases (GPx; EC 1.11.1.9 and EC 1.11.1.12) constitute a family of antioxidant enzymes widely found in animals, plants, fungi, and bacteria, and are implicated in protecting cells against damaging ROS (Margis et al., 2008; Trenz et al., 2021). GPx catalyzes the reduction of H_2_O_2_ and organic hydroperoxides using glutathione (GSH) as an electron donor, contributing to the maintenance of redox homeostasis (Brown et al., 2000; Margis et al., 2008). The presence of GPx-like proteins in bacteria has been considered widespread and even ancient. However, their sequences, phylogeny, structures, and metabolic roles have not been thoroughly investigated (Zhang et al., 2023). As a metabolically versatile soil bacterium, *P. putida* may encounter diverse environmental stresses, including oxidative challenges arising from redox reactions during lignin or aromatic compounds degradation. Glutathione metabolism is important as a first line of defense under oxidative stress (Akkaya et al., 2018; Nikel et al., 2021). Interestingly, an enzyme, from the glutathione-S-transferase superfamily, Ds-GST1, was recently characterized as a β-etherase capable of cleaving the β-O-4 aryl ether bond of a dimeric lignin model compound (Marinović et al., 2018). The glutathione system of *P. putida* has, however, not been fully explored as compared to other bacteria, including *Escherichia coli* (Smirnova & Oktyabrsky, 2005).

In this work, we aimed at the identification of enzymes specifically involved in lignin degradation by *P. putida* KT2440. Candidate enzymes were initially identified via comparative genome analysis and protein-BLAST. This surprisingly revealed putative involvement of a GPx-family enzyme in lignin metabolism by *P. putida*. Gene deletion, growth assays on lignin-derived substrates, dye decolorization assays, HPLC analysis, and transcriptomics were subsequently used to further corroborate a link between oxidative stress control and lignin-dependent metabolism in *P. putida*.

## 2. Materials and methods

### 2.1. Bacterial strains and growth conditions

Strains and plasmids are listed in Table 1. All *P. putida* strains were grown at 30°C in Lysogeny Broth (LB) medium containing 10 g L□^1^ tryptone, 5 g L□^1^ yeast extract and 5 g L□^1^ sodium chloride at 30°C with 200 rpm shaking. *Escherichia coli* strains were grown in LB at 37°C with 200 rpm shaking in horizontal shaker (Innova 4330, New Brunswick Scientific). For solid cultivation, 1.5 % (wt/vol) agar was added to LB. The M9 minimal medium used for growth assay was supplemented with 2 mg L□^1^ MgSO_4_ (Hartmans et al., 1990). M9 minimal medium containing either 10 mM CA, 10 mM FA, 10 mM SA, or a mixture of 5 mM CA, 1 mM FA, and 0.5 mM SA was used for degradation assay. When indicated, 50 µg mL□^1^ kanamycin, 30 µg mL□^1^ gentamycin, and 100 µg mL□^1^ streptomycin were added to the media.

**TABLE 1.**
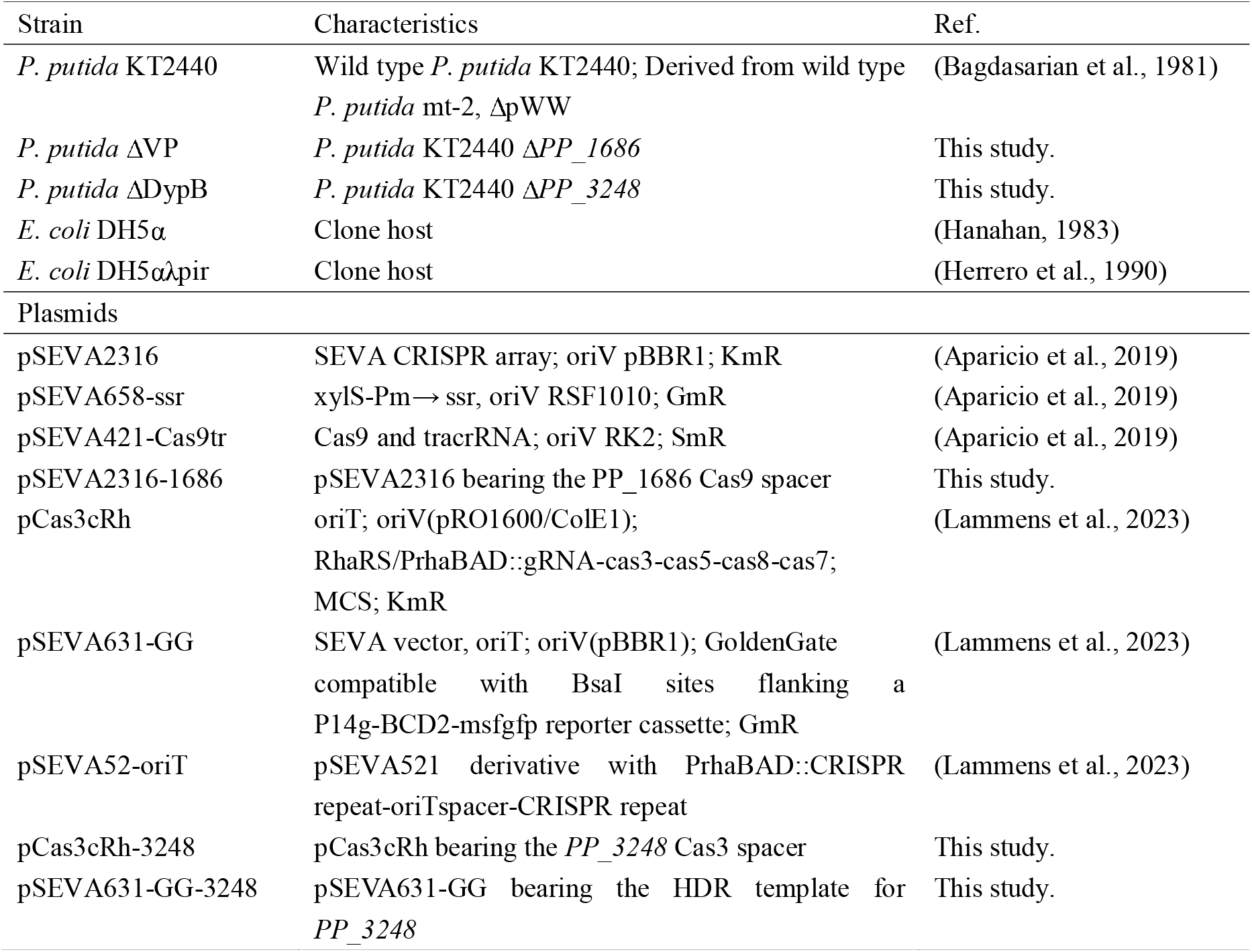
Strains and plasmids used in this study.

### 2.2 Enzyme BLAST and signal peptide prediction

Putative orthologous of lignin-degrading enzymes from *P. putida* NX-1 (Xu et al., 2018) were identified in *P. putida* KT2440 using the Basic Local Alignment Search Tool (BLAST) provided by NCBI (Altschul et al., 1990). To predict possible secretion signals, amino acid sequences of putative orthologous were analyzed using SignalP 6.0 (Nielsen et al., 2024).

### 2.3. Gene deletion and of *P. putida*

*P. putida* KT2440 genome with accession number AE015451.2 was used to target the genetic regions and determine primer sequences. Primers, spacers and ssDNA repair fragments used in this work are listed in Supplementary Table S1. Deletion of *PP_1686* was performed using CRISPR-Cas 9 as described previously (Aparicio et al., 2019). Deletion of *PP_3248* was performed using CRISPR-Cas3 3 as described previously (Lammens et al., 2023). Sanger sequencing was performed to confirm the gene manipulation.

PCR reactions were performed using Phire polymerase (Thermo Fischer) according to the manufacturer’s manual. Primers are listed in Table S1 and were obtained from Sigma-Aldrich. PCR products were visualized and analyzed by gel electrophoresis on 1% (w/v) TBE agarose gels containing 5 mg L^−1^ ethidium bromide in an electric field (110 V, 0.5× TBE running buffer).

### 2.4. Growth assays using lignin-derived aromatics

The *P. putida* strains were streaked from glycerol stocks onto LB agar plates and incubated for 12 h at 30 °C. Single colonies were inoculated into 5 ml of fresh LB medium and cultivated overnight at 30°C with shaking at 200 rpm. Overnight cultures were then inoculated into fresh M9 minimal medium supplemented with specific concentrations of *p*-coumaric acid, ferulic acid, syringic acid, or their mixture at an initial OD_600_ of 0.02. The molar ratio of *p*-coumaric acid, ferulic acid, and syringic acid in mixed substrates is 10:2:1, reflecting the ratio found in alkaline pretreated liquor from corn stover (Liu et al., 2024; Liu et al., 2017). Growth was monitored by measuring OD_600_ using a TECAN Spark 10M multimode microplate reader.

### 2.5. High-performance liquid chromatography (HPLC) analysis

*P. putida* strains were streaked from glycerol stocks onto LB agar plates and incubated for 12 h at 30 °C. Single colonies were inoculated into 10 ml of fresh LB medium and cultivated overnight at 30°C with shaking at 200 rpm. Overnight cultures were then inoculated into a 250-mL shake flask containing 50 mL M9 minimal medium supplemented with specific concentrations of *p*-coumaric acid, ferulic acid, syringic acid, or their mixture at an initial OD_600_ of 0.02. Samples were collected at 8, 12, 24, and 48 h. OD_600_ was measured to monitor the growth. For HPLC analysis, the samples were centrifuged, and the resulting supernatants were diluted 1:1 with acetonitrile to a final volume of 1 mL, followed by filtration through 0.2-µm syringe filters (Sartorius). The prepared samples were analyzed using a Shimadzu LC2030 HPLC system equipped with Shimadzu Shim-Pack GIST-HP C18-AQ column (3.0□×□150□mm, 3□μm) at 40 °C equipped with a UV detector monitoring at 240 and 254□nm. The mobile phases consisted of (A) 95/5 water/acetonitrile with 0.1% trifluoroacetic acid and (B) 5/95 water/acetonitrile with 0.1% trifluoroacetic acid, with the following gradient: 5% B for 0–2 min, 5–50% B from 2–9.5 min, 50–100% B from 9.5-10 min, 100% B for 10–12 min, 100–5% B from 12–13 min, and 5% B for 13–15 min.

The degradation percentage was calculated based on the reduction in substrate peak area over time, using the following equation:

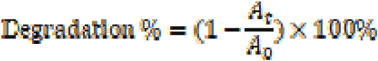

*A*_*0*_ represents the peak area at 0 h, and *A*_*t*_ represents the peak area at time *t*. All analyses were performed in triplicate.

### 2.6. Methylene blue decolorization assays

Lignin-like dye, methylene blue (MB), was used for ligninolytic activity assays. The decolorization ability of *P. putida* strains was determined using a quantitative dye decolorization assay, as described by Bharti et al (Bharti et al., 2019) with modifications. Overnight cultures were inoculated into fresh LB until the OD_600_ reached 1.0. Aliquots of 200 µL were transferred into 96-well plates supplied with 50 mg L^-1^ MB and incubated at 30 °C for 72 hours. Following incubation, the culture was centrifuged, and the absorbance of the supernatant was measured at 665 nm (λmax of MB) using TECAN Spark 10M multimode microplate reader. Cell-free incubations were assayed as control. The decolorization ratio was measured according to the following equation:

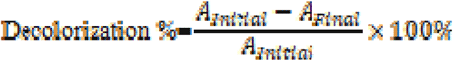

### 2.7. RNA-seq analysis

Wild-type and ΔVP strains of *P. putida* KT2440 were cultivated in M9 minimal medium supplemented with a mixture of 5 mM p-coumaric acid, 1 mM ferulic acid, and 0.5 mM syringic acid (10:2:1 molar ratio). Cell pellets were collected at 12 h and shipped to Novogene (Germany) for transcriptome analysis. Total RNA extraction, library preparation, and sequencing were performed by Novogene using standard protocols. The bioinformatic analysis was carried out using the Galaxy platform (http://usegalaxy.eu). Clean reads were mapped to the *P. putida* KT2440 reference genome with HISAT2, gene counts were obtained using Feature Counts, and differential expression analysis was performed with DESeq2. Genes with |log□ fold change| ≥ 1 and adjusted p < 0.05 were considered significantly differentially expressed. RNA-seq analysis was performed from three independent replicates for each sample.

## 3. Results and discussion

### 3.1. Enzymes BLAST and analysis

Genes in *P. putida* KT2440 encoding orthologous enzymes putatively involved in lignin decomposition in *P. putida* NX-1 (Xu et al., 2018) were identified using BLAST (Table 2). Since lignin is too large to be taken up by the cell, we particularly focus on putative extracellular enzymes (Table 2). SignalP6.0 was used to predict potentially secreted enzymes. Interestingly, although no signal peptide was predicted for the Dyp-type peroxidase in KT2440, previous secretome studies indicated their extracellular presence (Xu et al., 2022).

Surprisingly, a gene annotated in *P*. putida NX-1 to encode Versatile Peroxidase (VP) appeared similar to *PP_1686*, one of two genes annotated as glutathione peroxidase (GPx) in *P*. putida KT2440. Multiple sequence alignment between this NX-1 VP-homolog and both annotated KT2440 GPx genes, *PP_1686* and *PP_0777* revealed that *PP_1686* shared higher sequence similarity and coverage with the NX-1 homolog than *PP_0777* (Fig. 1). Notably, *PP_1686* retains most of the conserved residues present in the NX-1 sequence, particularly in the regions corresponding to the active site, whereas *PP_0777* shows several deletions and lower alignment continuity. In addition, both the NX-1 homolog and *PP_1686* contain an N-terminal hydrophobic stretch predicted as a TAT signal peptide, which is absent from *PP_0777*. The apparent absence of N-terminal part of NX-1 VP-homolog is most likely because of an annotation mistake (Xu et al., 2018). These observations indicate that *PP_1686* is a closer homolog of the NX-1 enzyme in *P. putida* KT2440. Both VP-homologues may represent candidates for investigating potential extracellular peroxidase activity. Despite these similarities, the physiological role of *PP_1686* and its NX-1 homologue in lignin degradation remains to be confirmed.

**Fig. 1.**
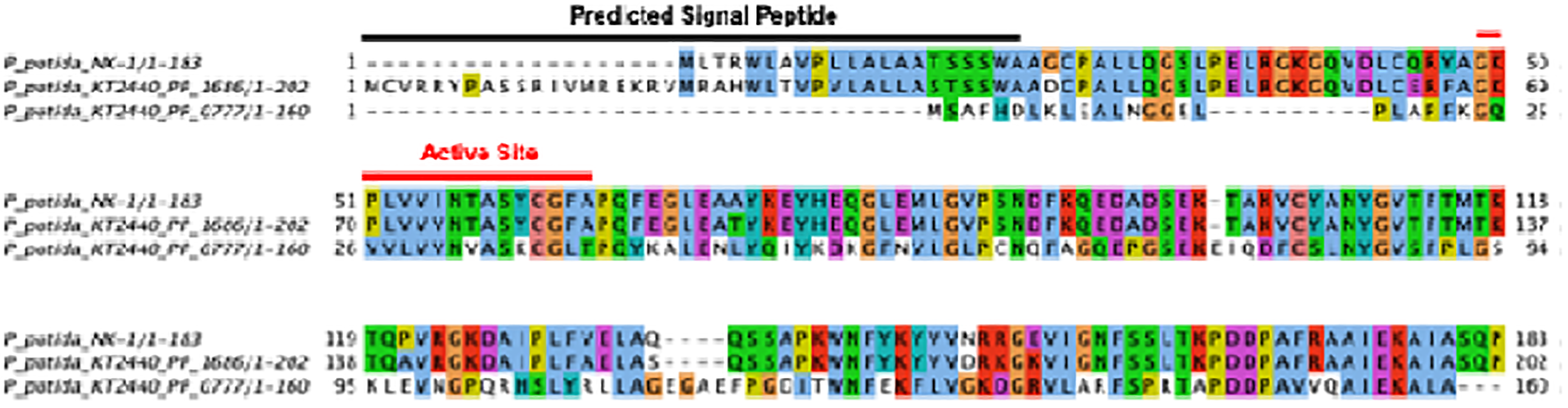
Multiple sequence alignment of glutathione peroxidase homologs from *P. putida* NX-1 and *P. putida* KT2440. Conserved residues are color-coded according to amino acid similarity. The predicted signal peptide and active site regions are annotated above the alignment.

**TABLE 2.**
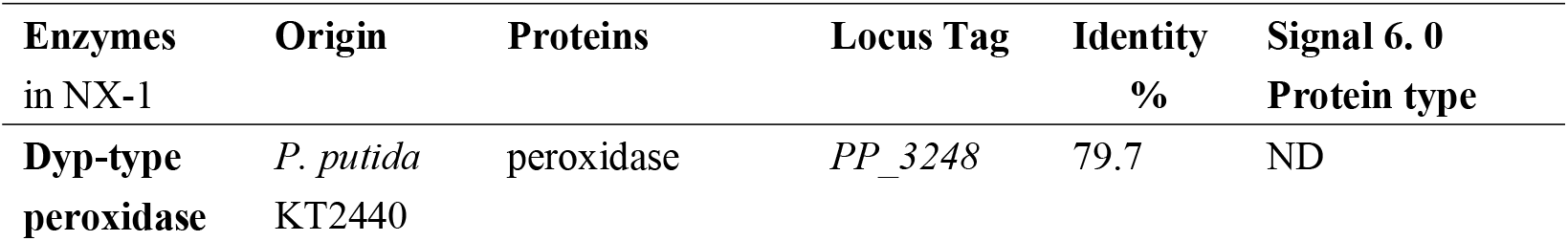
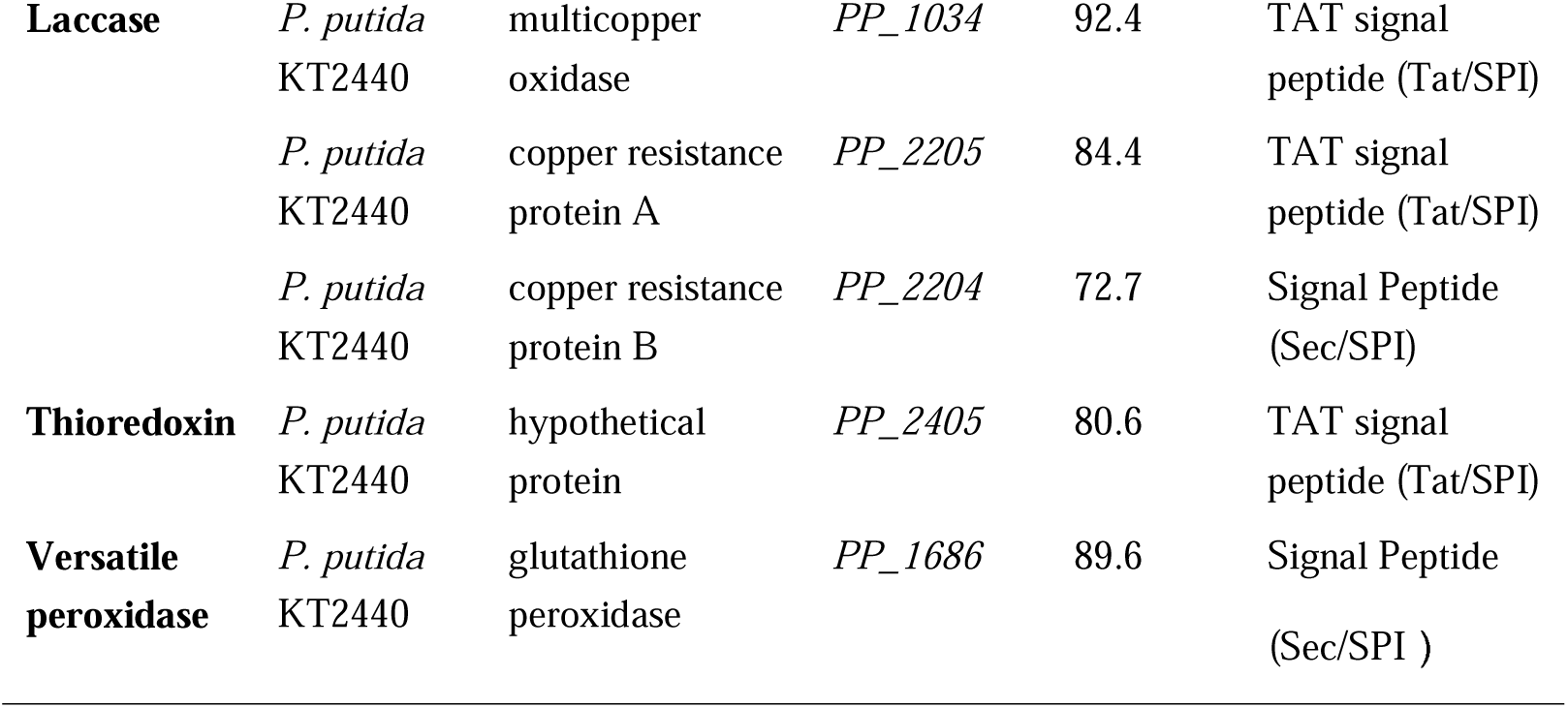
*P. putida* KT24 40 chromosome encodes enzymes orthologous to the enzymes implicated in lignin degradation in *P. putida* NX-1. ND, not detected.

Based on the BLAST analysis (Table 2), this study now focuses on characterization of the Versatile Peroxidase (VP) and Glutathione Peroxidase (GPx) homolog, *PP_1686*, and the Dyp-type peroxidase (DypB), *PP_3248. PP_1686* and *PP_3248* were deleted from the *P. putida* KT2440 genome by using CRISPR-Cas 9 and CRISPR-Cas 3 genome engineering approaches, resulting in the mutant strains *P. putida* ΔVP (ΔVP) and *P. putida* ΔDypB (ΔDypB). All constructs and deletion mutants were verified by PCR (Supplementary Fig. S1) and Sanger sequencing.

### 3.2. Growth on lignin-derived compounds

We started investigating the role of the *P. putida* VP/GPx and DypB enzymes by assessing growth of mutants on lignin-derived compounds (Fig. 2A-E). *p*-Coumaric acid (CA), ferulic acid (FA), and syringic acid (SA) were selected as representative model compounds of H-, G-, and S-lignin units, respectively (Liu et al., 2024; Notonier et al., 2021). In LB medium, wild-type KT2440 (WT), ΔVP, and ΔDypB did not show any growth differences, indicating that deletion of *PP_1686* or *PP_3248* does not impair growth under nutrient-rich conditions. When cultivated on CA, FA, or on a mixture medium, the WT strain displayed robust growth, whereas the ΔVP mutant showed reduced growth, particularly on CA. No significant growth difference was observed for the ΔDypB mutant. Importantly, all strains were unable to grow on SA.

**Fig. 2.**
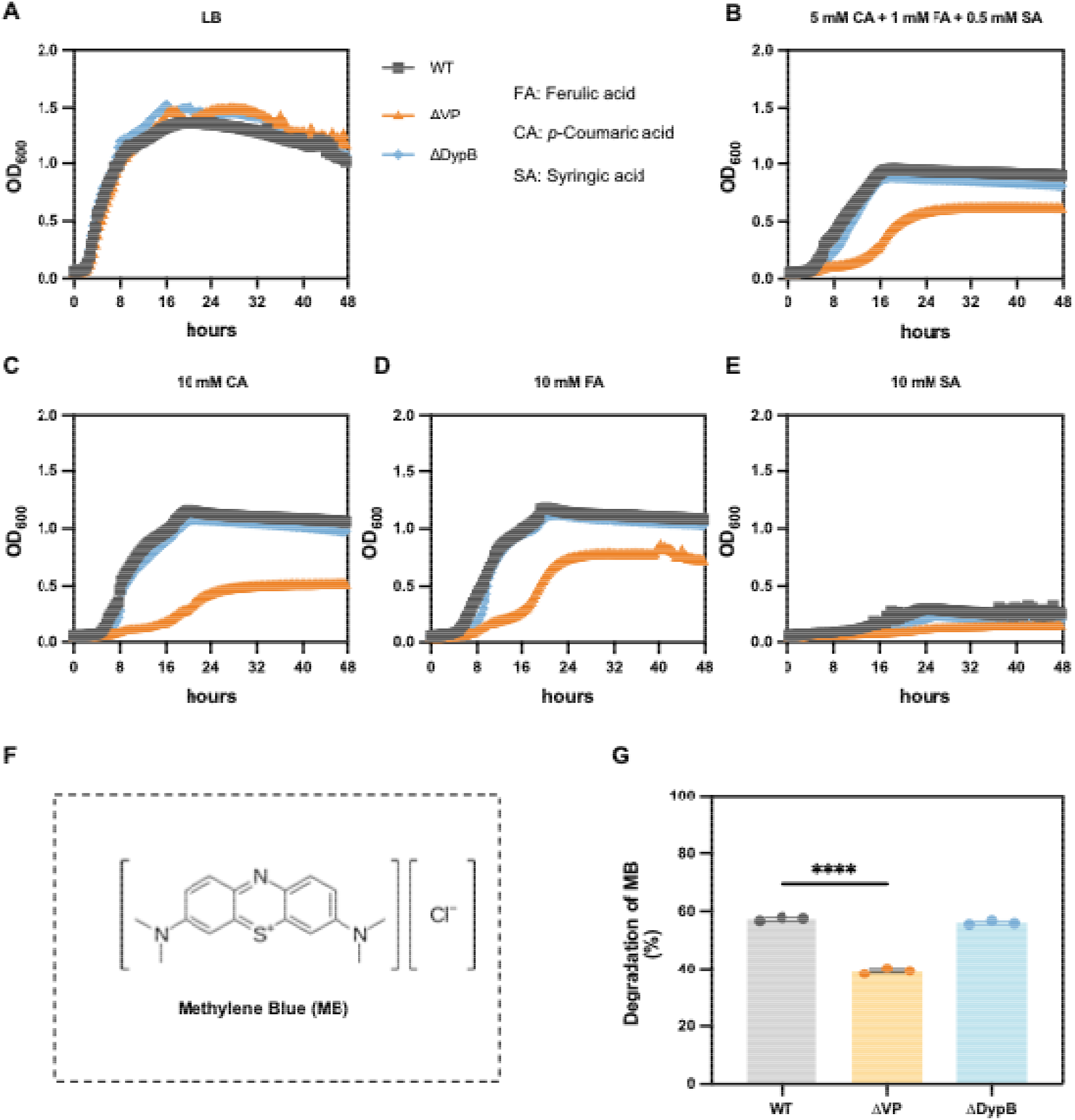
Growth of *P. putida* strains on lignin-derived compounds and degradation of methylene blue (MB). (A) Growth of WT, ΔVP, and ΔDypB in LB medium. (B-E) Growth of WT, ΔVP, and ΔDypB in M9 minimal medium supplemented with either a mixture of 5 mM *p*-coumaric acid, 1 mM ferulic acid, and 0.5 mM syringic acid, or 10 mM *p*-coumaric acid, 10 mM ferulic acid, 10 mM syringic acid as sole carbon sources, respectively. OD_600_ was measured over time using a microplate reader. (F) Structure of MB. (G) Quantitative MB degradation assay.

KT2440 is known to metabolize CA and FA through the β-ketoadipate pathway (Harwood & Parales, 1996; Jiménez et al., 2002; Nogales et al., 2017). Despite the presence of a putative native pathway for SA, KT2440 cannot grow on SA as the sole carbon source (Belda et al., 2016; Sonoki et al., 2018). Clearly, loss of *PP_1686* negatively affected the metabolism of CA and FA, indicating that this gene plays an important role in the utilization of these lignin-derived compounds.

### 3.3. Degradation of ligninolytic indicator dyes

To further investigate ligninolytic potential independently of lignin utilization, decolorization of synthetic lignin-mimicking dye methylene blue (MB, Fig. 2F) was monitored (Husain, 2006). Dye degradation was assessed for WT and mutant strains in liquid assays with growing cultures (Fig. 2G). The WT strain degraded MB by approximately 57%, whereas the ΔVP only degraded around 39%. By contrast, and importantly, no significant differences were observed between WT and ΔDypB.

These results indicate that *PP_1686* contributes to MB degradation, whereas *PP_3248* is dispensable under the tested conditions. The reduced degradation of ΔVP further demonstrates that *PP_1686* plays a functional role in metabolism of lignin-derived compounds, while *PP_3248* appears not essential under the tested conditions.

### 3.4. Role of *PP_1686* in lignin-derived compound utilization

To assess the role of *PP_1686* in the utilization of *p*-coumaric acid, growth experiments were repeated in shake flasks and supernatant analyzed by HPLC (Fig. 3-6).

**Fig. 3.**
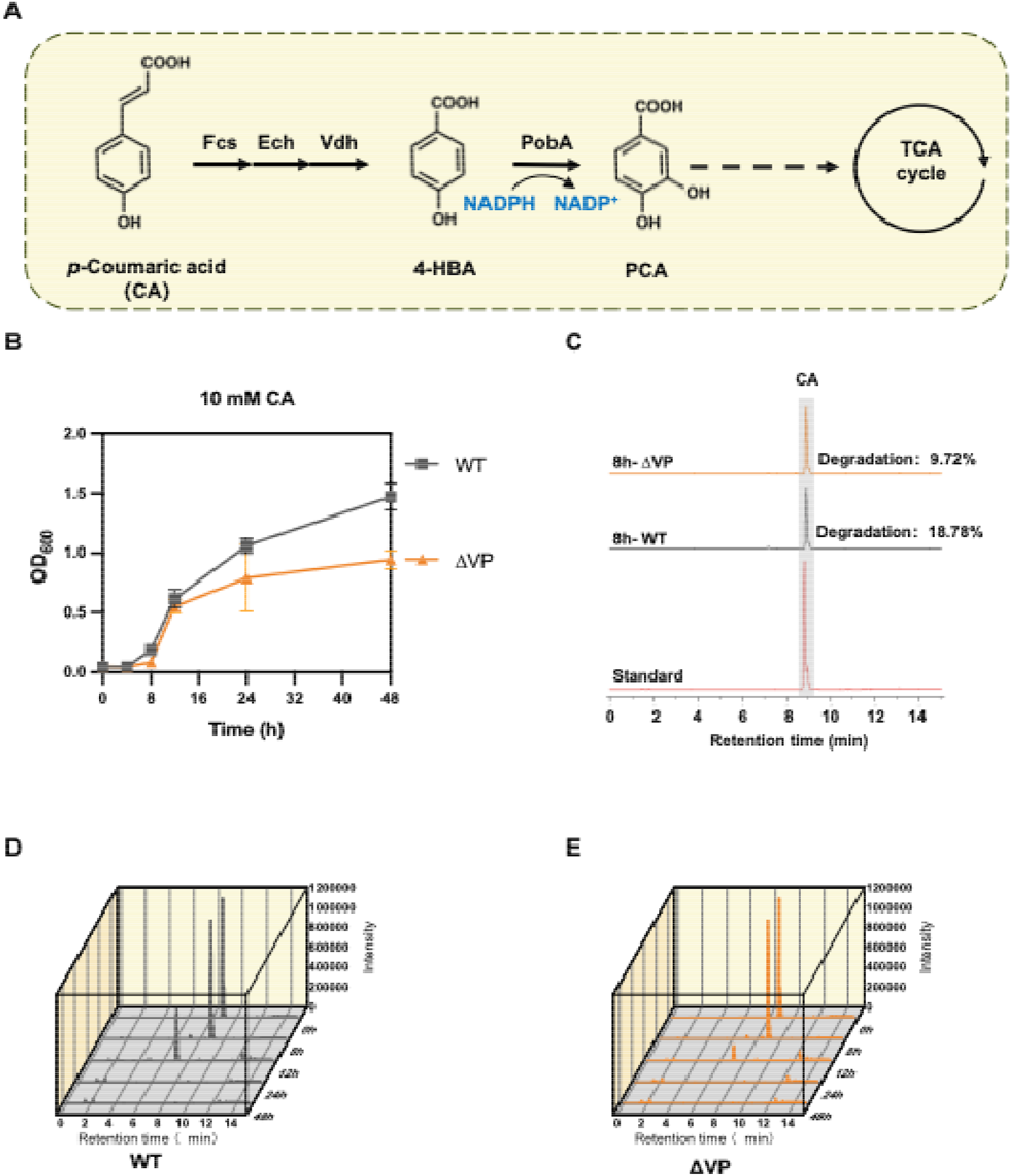
Role of PP_1686 in *p*-coumaric acid catabolism in *P. putida* KT2440. (A) *p*-coumaric acid (CA) degradation pathway in *P. putida* KT2440. (B) Growth of WT and ΔVP in M9 minimal medium supplemented with 10 mM CA as the sole carbon source. (C) Representative HPLC chromatograms at 8 h showing residual CA in WT and ΔVP strains compared with a CA standard. (D–E) Time-course HPLC analysis of CA consumption by WT (D), ΔVP (E) during 48 h of cultivation. Fcs, acyl-CoA synthetase; Ech, enoyl-CoA hydratase; Vdh, vanillin dehydrogenase; 4-HBA, 4-hydroxybenzoic acid; PobA, p-hydroxybenzoate hydroxylase; PCA, protocatechuic acid.

In *P. putida* KT2440, *p*-coumaric acid is first converted by acyl-CoA synthetase (Fcs), enoyl-CoA hydratase (Ech), and vanillin dehydrogenase (Vdh) into 4-hydroxybenzoic acid (4-HBA), which is subsequently hydroxylated by *p*-hydroxybenzoate hydroxylase (PobA) to protocatechuic acid (PCA) and further metabolized through the β-ketoadipate pathway into the TCA cycle (Fig. 2A). The WT strain exhibited robust growth on CA, reaching an OD_600_ of ~1.5 after 24 h. In contrast, the ΔVP mutant showed impaired growth.

To corroborate these results, we performed HPLC analysis of the culture supernatants. After 8 h of cultivation, residual CA was slightly higher in the ΔVP mutant compared to the wild type. Time-course analysis revealed that CA was efficiently consumed by both the wild-type and ΔVP strains. A slightly slower depletion rate in the ΔVP mutant corresponded with reduced accumulation of an intermediate peak with a retention time of ~7 min (Fig. 3D–F) suggesting a reduced CA conversion efficiency along the catabolic pathway. Although the identity of the intermediate was not determined in this study, its transient accumulation pattern may reflect a reduced conversion efficiency or slower turnover of *p*-coumaric acid degradation pathway intermediates. Despite the eventual depletion of CA in all strains, the ΔVP mutant showed lower OD_600_ values compared with the WT, indicating reduced biomass formation. These findings demonstrate that *PP_1686* contributes to optimizing aromatic catabolism of lignin-derived compounds, enhancing both metabolic efficiency and biomass formation, rather than being strictly essential for *p*-coumaric acid utilization.

Ferulic acid (FA) is also converted via Fcs, Ech, and Vdh to vanillic acid (VA) in *P. putida* KT2440, which is subsequently demethylated by vanillic acid O-demethylase oxygenase (VanAB) to PCA and funneled into the TCA cycle (Fig. 4A). Both the wild-type and ΔVP mutant strains were able to grow on FA, reaching OD□□□ values around 1.0 after 24 h (Fig. 4B), although biomass formation remained slightly lower than the wild type. HPLC analysis confirmed that FA was progressively consumed by both strains (Fig. 4C–E). The wild type showed efficient depletion of FA within 24–36 h, while the ΔVP mutant exhibited slower consumption over the first 8-12 h (Fig. 4C).

**Fig. 4.**
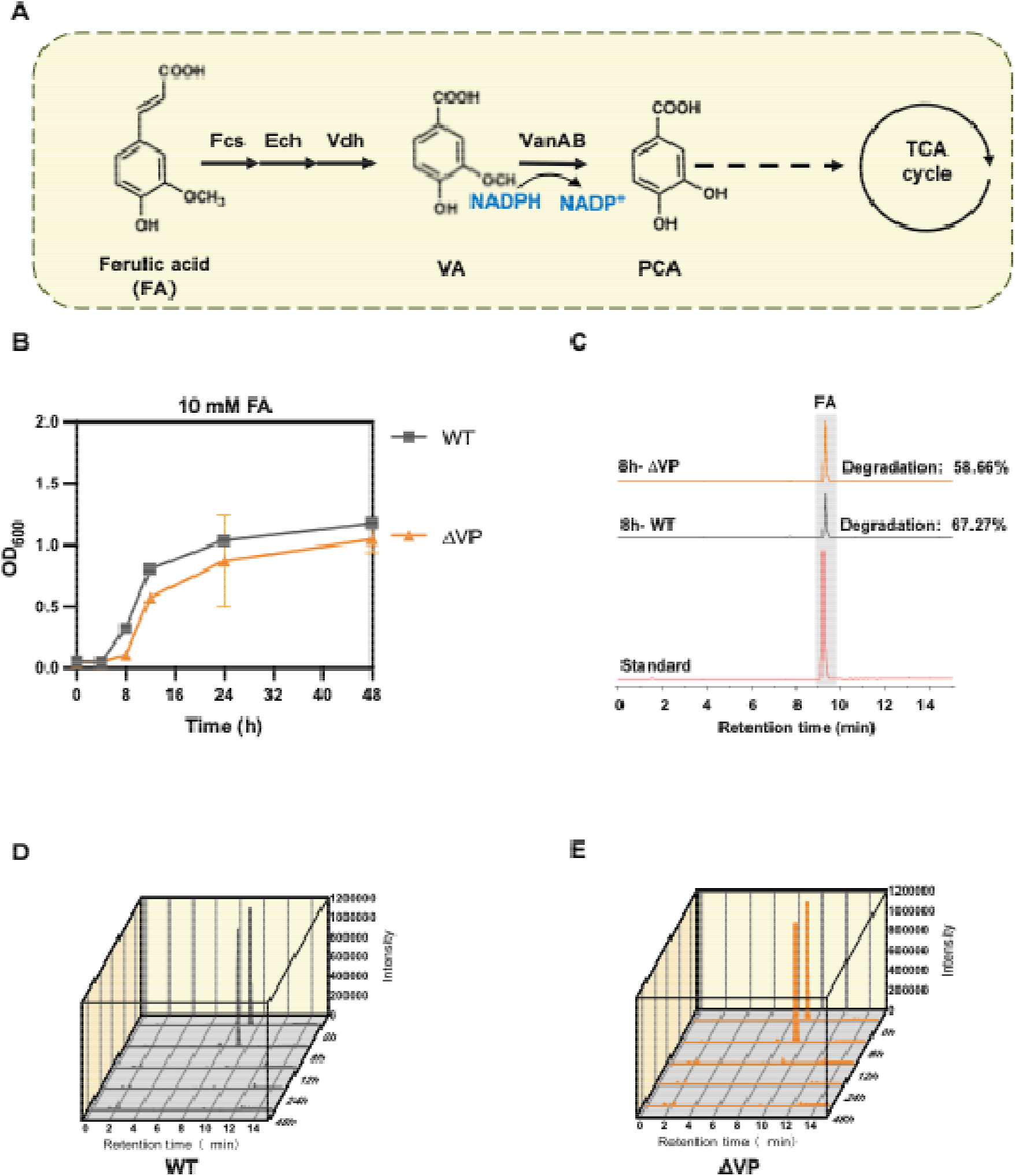
Role of PP_1686 in ferulic acid catabolism in *P. putida* KT2440. (A) Ferulic acid (FA) degradation pathway in *P. putida* KT2440. (B) Growth of WT and ΔVP in M9 minimal medium supplemented with 10 mM FA as the sole carbon source. (C) Representative HPLC chromatograms at 8 h showing residual FA in WT and ΔVP strains compared with a FA standard. (D–E) Time-course HPLC analysis of FA consumption by WT (D), ΔVP (E) during 48 h of cultivation. Fcs, acyl-CoA synthetase; Ech, enoyl-CoA hydratase; Vdh, vanillin dehydrogenase; VA, vanillic acid; VanAB, vanillic acid O-demethylase oxygenase; PCA, protocatechuic acid.

In contrast, no significant difference between P. putida KT2440 and the ΔVP strain in growth or substrate consumption was observed when 10 mM syringic acid (SA) was provided as the sole carbon source, consistent with previous reports that *P. putida* KT2440 is unable to metabolize S-type aromatics. The SA results are provided in Fig. S2.

To mimic more realistic lignin utilization conditions, a mixture of *p*-coumaric acid (5 mM), ferulic acid (1 mM), and syringic acid (0.5 mM) was tested as the sole carbon source. In mixed-substrate cultures, the WT grew faster and reached higher OD□□□ values compared with the ΔVP mutant, while the complemented strain largely restored this phenotype (Fig. 5A). HPLC analysis revealed that at 8 h, *p*-coumaric acid was consumed to a greater extent by the WT than by the ΔVP mutant (Fig. 5B). Ferulic acid was consumed progressively in all strains, although with slower metabolism in ΔVP compared to the WT. After 48h of cultivation, syringic acid was no longer detected under these conditions, suggesting that it may have been oxidized or converted through an alternative metabolic route rather than assimilated through the canonical pathway.

**Fig. 5.**
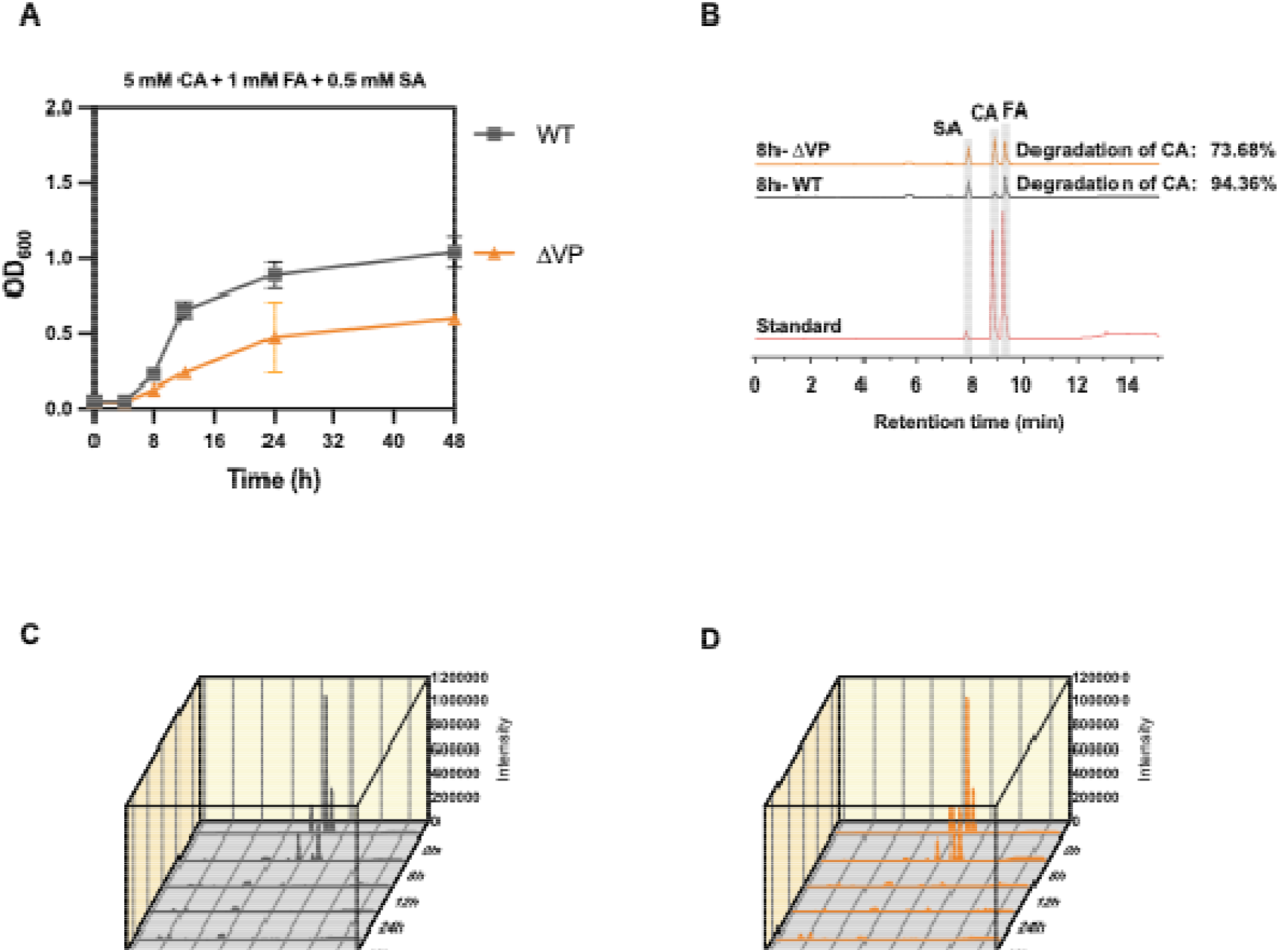
Role of PP_1686 in mixed lignin-derived compounds catabolism in *P. putida* KT2440. (A) Growth of WT and ΔVP in M9 minimal medium supplemented with 5 mM CA, 1 mM FA, and 0.5 mM SA. (B) Representative HPLC chromatograms at 8 h showing residual lignin-derived compounds in WT and ΔVP strains compared with a standard. (C–D) Time-course HPLC analysis of CA, FA and SA consumption by WT (C) and ΔVP (D) during 48 h of cultivation.

Our results show that *PP_1686* contributes to the efficient utilization of lignin-derived aromatics in *P. putida* KT2440. The effect of *PP_1686* deletion on aromatic catabolism was substrate dependent. In *p*-coumaric acid cultures, the ΔVP mutant showed markedly reduced growth and altered intermediate accumulation compared with the wild type. However, in ferulic acid cultures, the differences between WT and mutants were less pronounced. Although ΔVP exhibited slightly slower substrate consumption and reduced biomass formation relative to the wild type, both strains eventually depleted ferulic acid. These findings indicate that *PP_1686* contributes to the efficient utilization of multiple lignin-derived aromatics, with a stronger impact on *p*-coumaric acid metabolism.

### 3.5. Transcriptomic profiling confirms a role for *pobA* during growth on lignin-derived compounds

RNA-seq analysis was performed to compare the transcriptional profiles of the WT and ΔVP strains cultivated in mixed medium containing 5 mM *p*-coumaric acid, 1 mM ferulic acid, and 0.5 mM syringic acid for 12 h (Fig. 6). Among the differentially expressed genes, *pobA* (encoding 4-hydroxybenzoate hydroxylase), a key enzyme catalyzing the conversion of 4-hydroxybenzoate to protocatechuate in the β-ketoadipate pathway (Fig. 3A, Fig. 6A), was significantly down-regulated in ΔVP compared with the WT (Fig. 6B). This transcriptional change is consistent with the impaired consumption of *p*-coumaric acid observed in the mutant strain. During *p*-coumaric acid degradation, we observed that at 8 h, the ΔVP strain accumulated less of an intermediate than the WT, in line with down-regulation of *pobA* (Fig. 3). Furthermore, genes for ribosomal protein (*rpsT, rpsU, rpsP, rplU, rpmE, rpmB*), DNA recombination and repair, and stress response (*recA, recN*, and *dinB*) were found down regulated in ΔVP, suggesting a reduced capacity for coping with genotoxic or oxidative stress (Fig. 6D). Downregulation of *glmS* and *tpiA* further indicated a reduction in amino sugar and glycolytic flux.

**Fig. 6.**
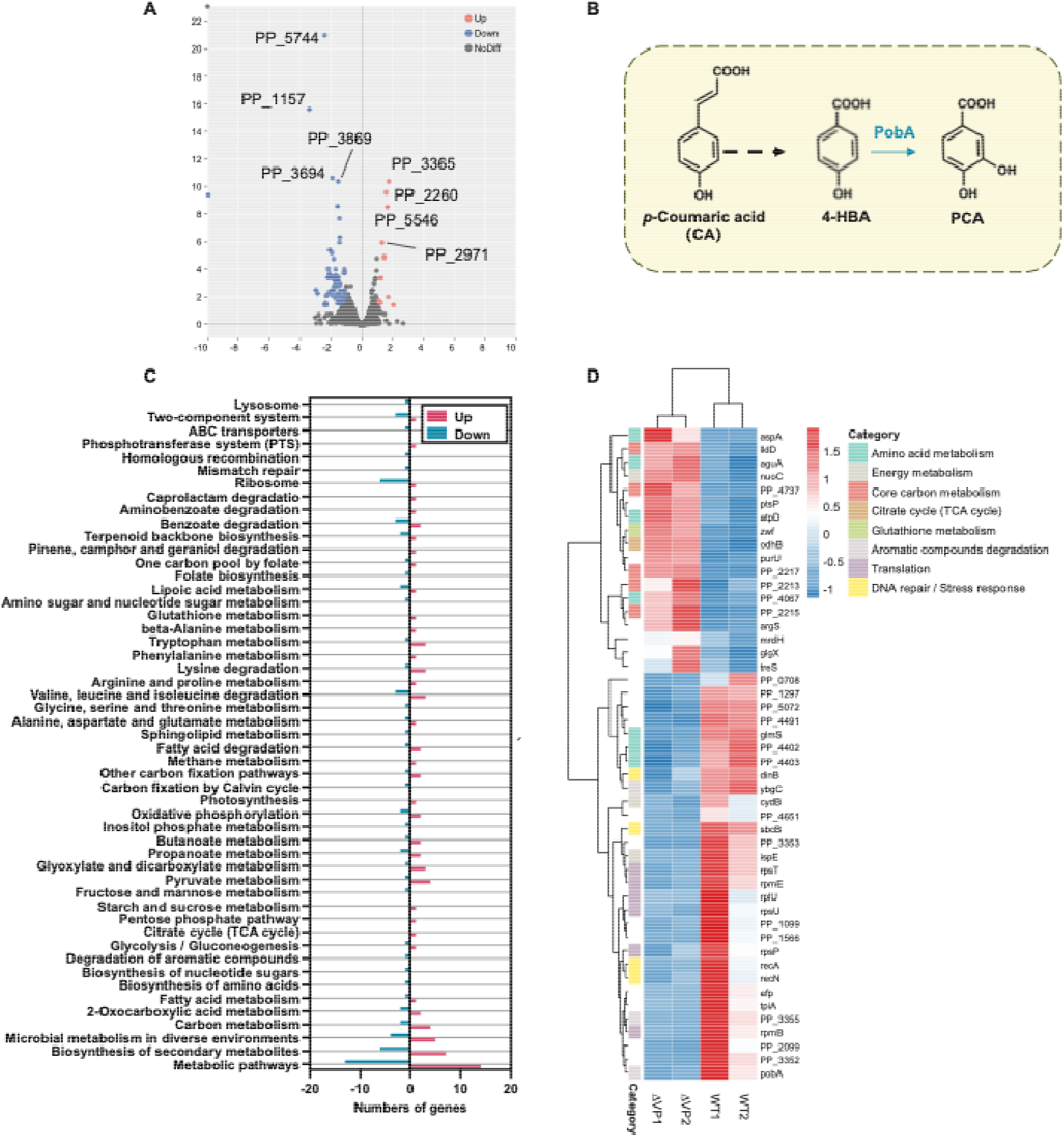
Transcriptomic analysis of WT and ΔVP strains cultured on mixed lignin-derived substrates. (A)Volcano plot showing differentially expressed genes (DEGs) between WT and ΔVP strains after 12 h in mixed medium (5 mM p-coumaric acid, 1 mM ferulic acid, 0.5 mM syringic acid). Blue, downregulated genes; red, upregulated genes; grey, not significant. Selected DEGs are highlighted. (B) Schematic of the *pobA* mediated *p*-coumaric acid catabolic pathway in *P. putida*. (C) KEGG pathway enrichment analysis of DEGs, showing the top significantly enriched metabolic pathways. Red bars indicate upregulated pathways; blue bars indicate downregulated pathways. (D) Targeted heatmap of selected DEGs grouped by functional categories, including amino acid metabolism, energy metabolism, core carbon metabolism, citrate cycle, glutathione metabolism, aromatic compound degradation, translation, and DNA repair/stress response. Expression values are shown as Z-scores of log□ fold change. Data represent three biological replicates for each condition.

In contrast, the ΔVP mutant exhibited transcriptional upregulation of several core metabolic pathways. Genes associated with the tricarboxylic acid (TCA) cycle and oxidative phosphorylation (*odhB, nuoC, atpD*) were induced, indicating a shift toward increased energy production and homeostasis. The pentose phosphate pathway gene *zwf* (glucose-6-phosphate dehydrogenase) was also upregulated, likely enhancing NADPH generation to support redox homeostasis in the absence of *PP_1686* (Fig. 6C-D). Overall, these changes suggest a compensatory reprogramming of central metabolism, enabling the ΔVP strain to redirect resources toward retrieving energy and redox balancing, causing impaired aromatic degradation.

Based on these results, we may further hypothesize on a regulatory role of *PP_1686*/VP in the utilization of lignin-derived compounds. Deletion of *PP_1686* directly resulted in the downregulation of *pobA*, while apparently impairing DNA repair and translational capacity. Remarkably, the mutant compensated for these deficiencies by enhancing TCA cycle activity, ATP synthesis, NADPH generation, and carbohydrate metabolism, reflecting re-establishment of the metabolic balance between catabolic efficiency and cellular homeostasis. These findings highlight a role of *PP_1686*/VP in coupling oxidative regulation and homeostasis with control of aromatic compound catabolism in *P. putida*.

## 4. Conclusions

Glutathione peroxidase (GPx) is regarded as one of the main antioxidant enzymes, which can reduce peroxides to compounds with lower toxicity (Margis et al., 2008). GPxs have been extensively studied and well-classified in mammals, where their roles in peroxide detoxification and redox regulation are well established (Bao et al., 2023; Trenz et al., 2021). In contrast, studies in bacteria remain scarce, and the physiological roles, evolutionary diversification, and functional relevance of bacterial GPxs are still poorly understood (Bao et al., 2023).

The Versatile Peroxidase/Glutathione Peroxidase, VP/GPx knock-out mutant in this study showed impaired growth and degradation of lignin-derived aromatics and a lignin-model dye compound in *P. putida* KT2440. Transcriptome analysis revealed that the loss of VP/GPx triggered a coordinated transcriptional reprogramming, including repression of key aromatic funneling genes (e.g., pobA), downregulation of DNA repair and translational modules, and upregulation of energy-generating pathways (e.g., nuo, atp, odhB), NADPH supply routes (e.g., zwf), and carbohydrate metabolism (e.g., treS, glgX). These findings suggest that VP/GPx functions in regulating oxidative metabolic balance in *P. putida*. Its loss uncouples oxidative defense from respiration and carbon flux, leading to concurrent deficits in aromatic metabolism and growth. Beyond its annotated role in ROS detoxification, our findings demonstrate that the VP/GPx glutathione peroxidase encoded by *PP_1686* has another role in supporting lignin degradation and aromatic catabolic fitness in *P. putida*. Together, these results point out a redox-dependent growth-coupling mechanism linking oxidative stress regulation with lignin degradation and utilization.

*P. putida* is emerging as a highly promising microbial chassis for lignin valorization, especially for its metabolic versatility and its recently revealed auxiliary enzyme network that enhances lignin degradation ability (Liang et al., 2025). A deeper understanding of its redox regulation and stress-related mechanisms is essential for rationally engineering robust and efficient lignin-degrading cell factories. This study broadens the understanding of bacterial GPxs, revealing an important contribution to the integration of oxidative stress response with carbon metabolism. Beyond annotated ROS detoxification, GPx appears as an auxiliary enzyme to play a central role in aligning redox balance with lignin degradation in *P. putida*. Moreover, our transcriptomic data uncovered a cluster of genes with hitherto unknown functions that were consistently downregulated in the VP/GPx knock-out mutant. Characterization of those genes may uncover novel proteins or pathways linked to lignin catabolism and stress adaptation. Future work should aim to dissect the molecular mechanisms by which GPx influences regulatory networks. Such work will not only enrich our understanding of redox metabolism coupling but also provide novel insights for the engineering of robust microbial platforms for lignin valorization.

## Notes

### Competing Interest Statement

The authors have declared no competing interest.

### Funding

- China Scholarship Council

