## Supplemental data for "Lignin degradation and valorization by *Pseudomonas putida* KT2440: a new role for glutathione peroxidase"

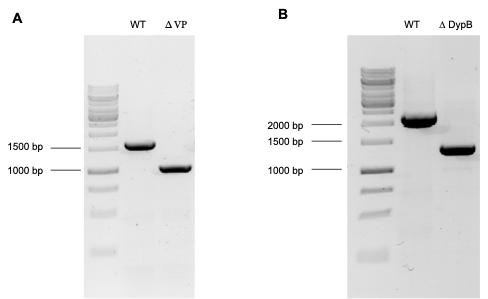
**Fig S1** PCR verification of gene deletion and complementation mutants. (A) PCR analysis of PP_1686 mutants. WT: *P. putida* KT2440 wild type; ΔVP: *P. putida* KT2440 PP_1686 deletion mutant; PP_1686: 609 bp. Primers: Seq-1686 F/ Seq-1686 R. (B) PCR analysis of PP_3248 mutants. WT: *P. putida* KT2440 wild type; ΔVP: *P. putida* KT2440 PP_3248 deletion mutant; PP_3248: 864 bp. DNA size markers are indicated on the left. Primers: Seq-3248 F/ Seq-3248 R.

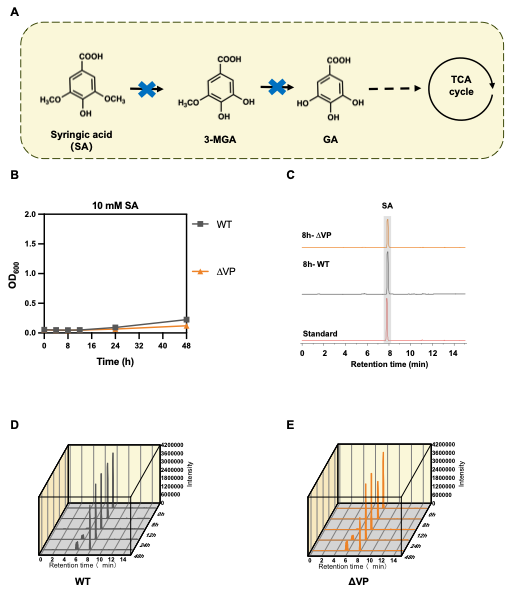
**Fig S2** Role of *PP_1686* in mixed lignin-derived compounds catabolism in *P. putida* KT2440. (A) Growth of WT, ΔVP in M9 minimal medium supplemented with 10 mM SA. (B) Representative HPLC chromatograms at 8 h showing residual lignin-derived compounds in WT and ΔVP strains compared with a standard. (C–E) Time-course HPLC analysis of SA consumption by WT (C), ΔVP (D) during 48 h of cultivation.

| **Oligos** | **Sequence** |
| --- | --- |
| pSEVA2316-F | TTAATTAAAGCGGATAACAATTTCACACAGGAG |
| pSEVA2316-R | ACTAGTCGCCAGGGTTTTCC |
| PP1686-S | AAACCACCATGACCAAGACACAGGCGGG |
| PP1686-AS | GTGGTACTGGTTCTGTGTCCGCCCAAAA |
| PP_1686-ssDNA | TGGCATCAGGGTGGGATGGCGCTGATGCAACTGGGCATGACC  AGGCTGCGCTTGCTGCGGCGTTGCTTTAAAGCGCGAGG |
| pCas3cR-F | CAGGAAATGCGGTGAGC |
| pCas3cR-R | GAGCAGCTAATTCACCGC |
| SEVA PS1 | AGGGCGGCGGATTTGTCC |
| SEVA PS2 | GCGGCAACCGAGCGTTC |
| PP_3248-S | GAAACCAGCAAGGTCTGCTTGCCACCCCGGTGCCGGCCCG |
| PP_3248-AS | GCGACGGGCCGGCACCGGGGTGGCAAGCAGACCTTGCTGG |
| PP_3248-U-F | ATAGGTCTCACTAGGTGAGTTGACCGCCGTCAGGG |
| PP_3248-U-R | ATCTGGATGTTTAGAGAATCAGGCTAATCCTAAGGTAAAAACGGA |
| PP_3248-D-F | CTTAGGATTAGCCTGATTCTCTAAACATCCAGATACAGAAACACCCG |
| PP_3248-D-R | ATAGGTCTCAGACTCAGCAAGTCGACCTTGTCGG |
| Seq-3248 F | CGGTTTTTCCATGCGCGC |
| Seq-3248 R | ATACGGGCGACTTCGTCG |
| Seq-1686-F | TGGGCATGACCAGGCTGCGCT |
| Seq-1686-R | CCCTGTTATCCCTAGAAGCTTGCATGCCTGCAGGTCGACTTCGCATG  AGCAATCTGTCCCG |

**TABLE S1** Primers, spacers, and ssDNA repair fragments used in this study.
